# MinE recruits, stabilizes, releases, and inhibits MinD interactions with membrane to drive oscillation

**DOI:** 10.1101/109637

**Authors:** Anthony G. Vecchiarelli, Min Li, Michiyo Mizuuchi, Vassili Ivanov, Kiyoshi Mizuuchi

## Abstract

The MinD and MinE proteins of *Escherichia coli* self-organize into a standing-wave oscillator on the membrane to help align division at mid-cell. When unleashed from cellular confines, we find that MinD and MinE form a wide spectrum of patterns on artificial bilayers - static amoebas, traveling waves, traveling mushrooms, and bursts with standing-wave dynamics. We recently focused our cell-free studies on bursts because their dynamics closely resemble those found *in vivo*. The data unveiled a patterning mechanism largely governed by MinE regulation of MinD interaction with membrane. We proposed that the MinD to MinE ratio on the membrane acts as a toggle switch between MinE-stimulated recruitment or release of MinD from the membrane. Here we provide data that further refines and extends our model that explains the remarkable spectrum of patterns supported by these two ‘simple’ proteins.

## INTRODUCTION

The MinCDE system of *Escherichia coli* forms a cell-pole to cell-pole standing wave oscillator that prevents cell division near the cell poles (Hu and Lutkenhaus, 1999; Lutkenhaus, 2007; Raskin and de Boer, 1999a). MinD is an ATPase that, when bound to ATP, can dimerize and bind membrane via its membrane targeting sequence (MTS) (Hu and Lutkenhaus, 2003; Szeto et al., 2002; Wu et al., 2011; Zhou and Lutkenhaus, 2003). MinE also functions as a dimer and has MTSs that are considered to be ‘inactive’ while MinE is in solution (Ghasriani et al., 2010; Kang et al., 2010; Park et al., 2011). In its active state, MinE stimulates the ATPase activity of membrane-bound MinD (Hsieh et al., 2010; Hu et al., 2002; Vecchiarelli et al., 2016). MinE-stimulated ATP hydrolysis by MinD is likely coupled to MinD release from membrane (Vecchiarelli et al., 2016). The third and final component is the inhibitor of divisome assembly called MinC. MinC is a passenger protein on MinD that links MinD distribution on the membrane to divisome positioning. But MinC itself is not required for MinD/E oscillation (Hu and Lutkenhaus, 2000; Raskin and de Boer, 1999b). The perpetual chase and release of MinD by MinE on the membrane produces a time-averaged concentration of MinC that is lowest at mid-cell (Fu et al., 2001; Hale et al., 2001; Hu and Lutkenhaus, 1999; Meinhardt and de Boer, 2001; Raskin and de Boer, 1999b). The oscillation therefore promotes cell division at mid-cell by inhibiting division near the poles (Lutkenhaus, 2007). The remarkable oscillatory dynamics of this self-organizing system were first reported more than 15 years ago (Raskin and de Boer, 1999a), but the molecular mechanism remains enigmatic.

The Schwille and Mizuuchi groups, and very recently the Dekker group, have reconstituted Min patterning dynamics on supported lipid bilayers (SLBs) of varying lipid compositions (Ivanov and Mizuuchi, 2010; Loose et al., 2008; Vecchiarelli et al., 2016; Vecchiarelli et al., 2014; Zieske and Schwille, 2014) and under different confinement geometries (Schweizer et al., 2012; Zieske et al., 2016; Zieske et al., 2014; Zieske and Schwille, 2013; Zieske and Schwille, 2014; Caspi and Dekker, 2016) to elucidate the molecular mechanism governing oscillation *in vivo*. Propagating waves of MinD chased by MinE was the first type of pattern to be reconstituted on the bottom of an SLB-coated well (Loose et al., 2008). In our SLB-coated flowcell, the Min system formed a variety of patterns (Ivanov and Mizuuchi, 2010; Vecchiarelli et al., 2016). Under constant flow, a near spatially homogeneous oscillation was generated, where large swaths of the SLB were bound and released by MinD and MinE (Ivanov and Mizuuchi, 2010; Vecchiarelli et al., 2014). Stopping the flow resulted in a pattern spectrum, where the MinD and MinE density on the SLB determined the mode of patterning – amoebas, waves, mushrooms or bursts (Figure 1) (Vecchiarelli et al., 2016). At very high protein densities, MinD and MinE formed amoebas - circular MinD binding zones of uniform size that were stably surrounded by an E-ring (Ivanov and Mizuuchi, 2010; Vecchiarelli et al., 2016). At moderate densities, MinD and MinE self-organized into the travelling waves described above. The protein densities within amoebas or waves are far in excess of what is possible *in vivo*. Also, these patterns lack standing wave dynamics with nodes where the time-averaged local MinD concentration is minimum, as observed at mid-cell *in vivo*. Thus, it was difficult to decipher the mechanistic principles underlying these dissimilar patterns and how they relate to standing-wave oscillations *in vivo*.

**Figure 1.**
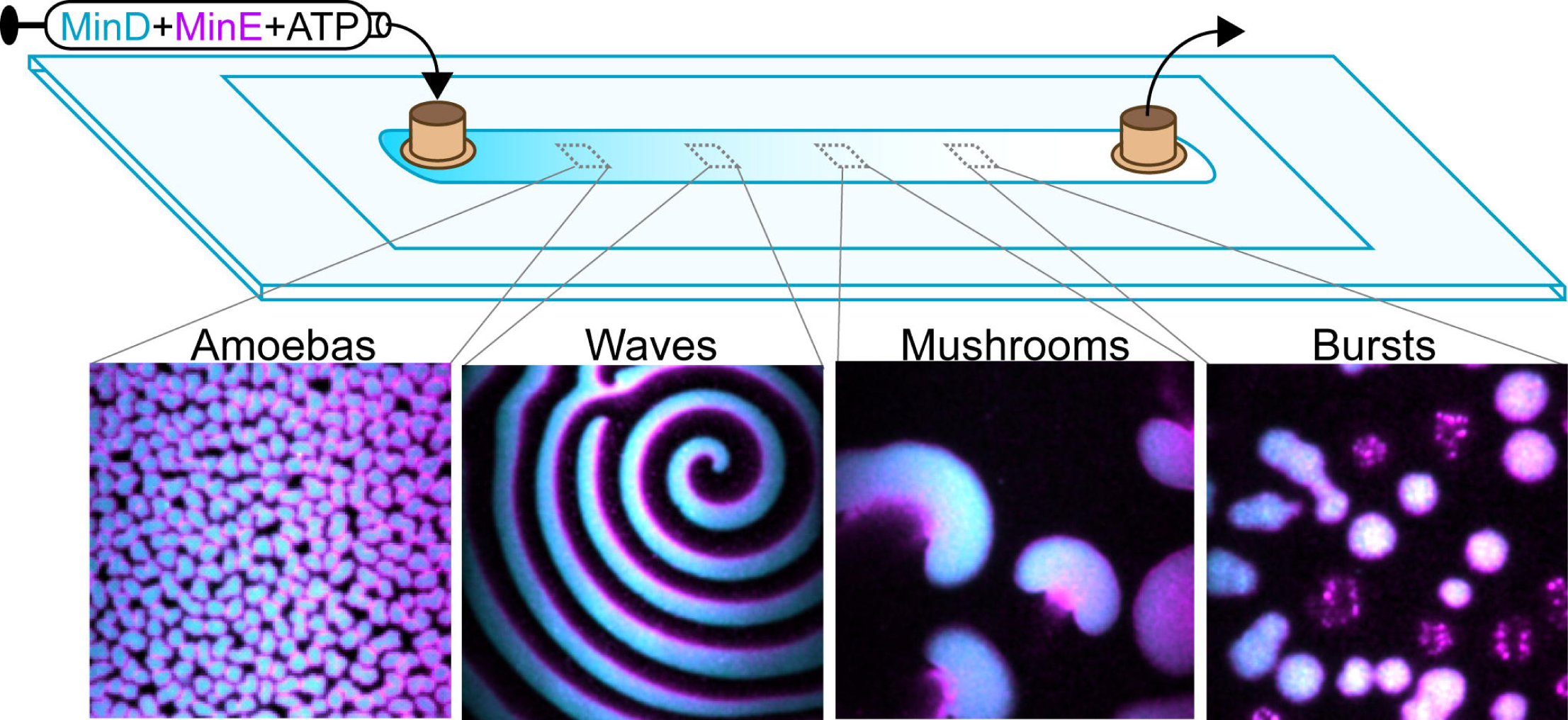
**MinD and MinE from a spectrum of cell-free patterns on a flat bilayer.** (**A**) GFP-MinD (cyan) and MinE-Alexa647 (magenta) were pre-incubated with ATP and infused for 10 minutes at 1 μl/min into a flowcell coated a supported lipid bilayer (SLB). Still images show the different modes of patterning supported by the decreasing GFP-MinD density on the SLB from inlet to outlet.

We recently used our flowcell setup to specifically address the mechanistic basis for standing-wave oscillations. We hypothesized that to reconstitute a standing-wave, the MinD supply must be limiting because when a MinD polar zone develops *in vivo* the cytoplasmic pool of MinD presumably depletes (Meinhardt and de Boer, 2001). Indeed, under protein depletion conditions, we observed two previously unidentified patterns we called mushrooms and bursts (Figure 1). Out of all patterns reconstituted on a flat SLB to date, only bursts displayed standing-wave dynamics. Bursts are radially expanding binding zones of MinD and MinE that initiated from a nucleation point on the SLB. MinD binding was rapid whereas MinE slowly accumulated within the MinD zone. As the local solution supply of MinD depleted, burst expansion halted, its perimeter was corralled by an E-ring, and the burst imploded (Vecchiarelli et al., 2016).

Mushrooms were an intermediate pattern between travelling waves and bursts where the MinD supply was semi-depleted (Figure 1). Like bursts, mushrooms temporally oscillated as expanding binding zones of MinD that were corralled and disassembled by MinE. In contrast to individual bursts, which were symmetric and spatially disconnected from one another, mushrooms budded out from the previous disassembling set of mushrooms, resulting in asymmetric propagation of the MinD binding zone, which was followed by a spatially skewed disassembly by MinE. After the MinD binding front of a mushroom stalled and was corralled by an E-ring, the spatial asymmetry was propagated by subsequent mushrooms. As the density of Min proteins increased on the SLB, mushrooms merged to form the propagating waves. This recent study focused on the burst pattern because it most closely resembled the standing-wave dynamics observed *in vivo*. The findings allowed us to propose a comprehensive molecular mechanism for standing-wave oscillations (Vecchiarelli et al., 2016). Here we provide data that further extends our explanation of how MinD and MinE support the wide variety of patterns formed both in and out of the cell.

## The Model

We propose that MinD and MinE dimers can independently and dynamically interact with membrane before any patterning event is even initiated (Figure 2A). Local fluctuations in the MinD to MinE ratio on the membrane are needed to nucleate the formation of a radially expanding binding zone containing both MinD dimers alone (D2) and those in complex with, and stabilized by, MinE (D2E2) (Figure 2B). D2E2 not only stabilizes MinD on the membrane, but also acts to rapidly recruit more MinD from solution *in vitro*, or from the cytoplasm *in vivo* (Figure 2C). Exactly how this recruitment complex, D2E2D2, plays a role in the patterning mechanism remains to be determined. MinE accumulates more slowly than MinD. Therefore, during pattern initiation the majority of MinE dimers are on the membrane in the D2E2 or D2E2D2 complex (Figure 2D). But as MinD binding slows down for any reason, such as surface exclusion or solution depletion, MinE binding will catch up and eventually tip the balance, where another MinE dimer can join a D2E2 complex and form E2D2E2 – the MinD dissociation complex (Figure 2E). Formation of this complex triggers ATP hydrolysis by MinD and its dissociation from the membrane. Therefore, in our model, the membrane-bound stoichiometry of MinD and MinE acts as the ‘switch’ from MinE-stimulated recruitment and stabilization of MinD on the membrane to MinE-stimulated release of MinD (Vecchiarelli et al., 2016).

**Figure 2.**
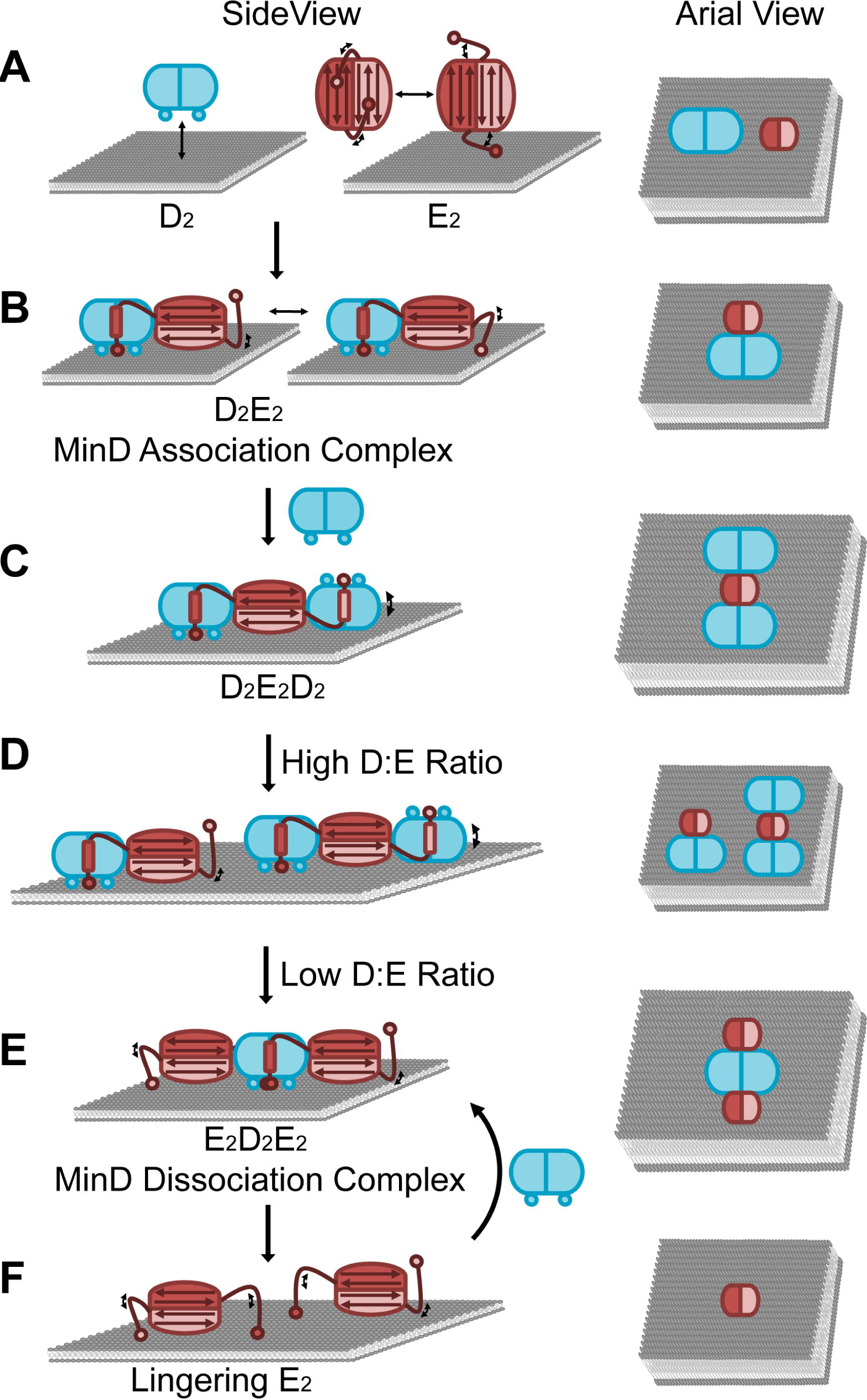
**MinD/MinE stoichiometry on the membrane drive patterning.** (**A**) MinD dimers (cyan) dynamically bind membrane. The closed form of MinE has a six-stranded β-sheet making its hydrophobic core. Conformational breathing of the MinE dimer allows for dynamic membrane binding via its MTSs (circles). (**B**) D2E2 complex formation stabilizes membrane association for both proteins. One MinD binding site is still available on the MinE dimer for interaction with another MinD dimer. (**C**) This second MinD binding site recruits more MinD from solution. (**D**) During patterning initiation, all MinE dimers on the membrane in complex as D2E2 or D2E2D2. (**E**) As MinE accumulates on the membrane, E2D2E2 complexes can form, which stimulate MinD release from the membrane. (**F**) MinE dimers linger on the membrane after MinD release, preventing MinD from rebinding the membrane. Arial View (right) highlights the difference in footprint size among the protein species.

When a MinD dimer is released, the MinE dimers responsible for its release can linger on the membrane (Figure 2F). This lingering MinE can then associate with other D2E2 complexes in the surrounding area, releasing more MinD and generating more lingering MinE dimers. At this high density of lingering MinE, any MinD dimers from solution, or diffusing from neighboring areas of the membrane, would be quickly joined by not one but two MinE dimers, and form the E2D2E2 complex (Figure 2E-F). This prevents MinD from re-accumulating on regions of the membrane that other MinD dimers have just dissociated from.

Let’s recap by taking the molecular mechanism back into the cell. A high density of lingering MinE provides the refractory period for MinD rebinding at the cell-pole from which it just dissociated (Figure 3A). Once the lingering MinE density sufficiently declines, they are joined by MinD dimers in a one-to-one complex, E2D2, which stimulates the recruitment of more MinD. The resulting D2E2D2 complex may then separate into D2E2 and D2 to continue the positive feedback cycle (Figure 3B-C). As the MinD polar zone grows, MinD depletes from the cytoplasm and MinE accumulates on the membrane. E2D2E2 then stimulates the release of MinD and the lingering MinE dimers concentrate to form an E-ring (Figure 3D).

**Figure 3.**
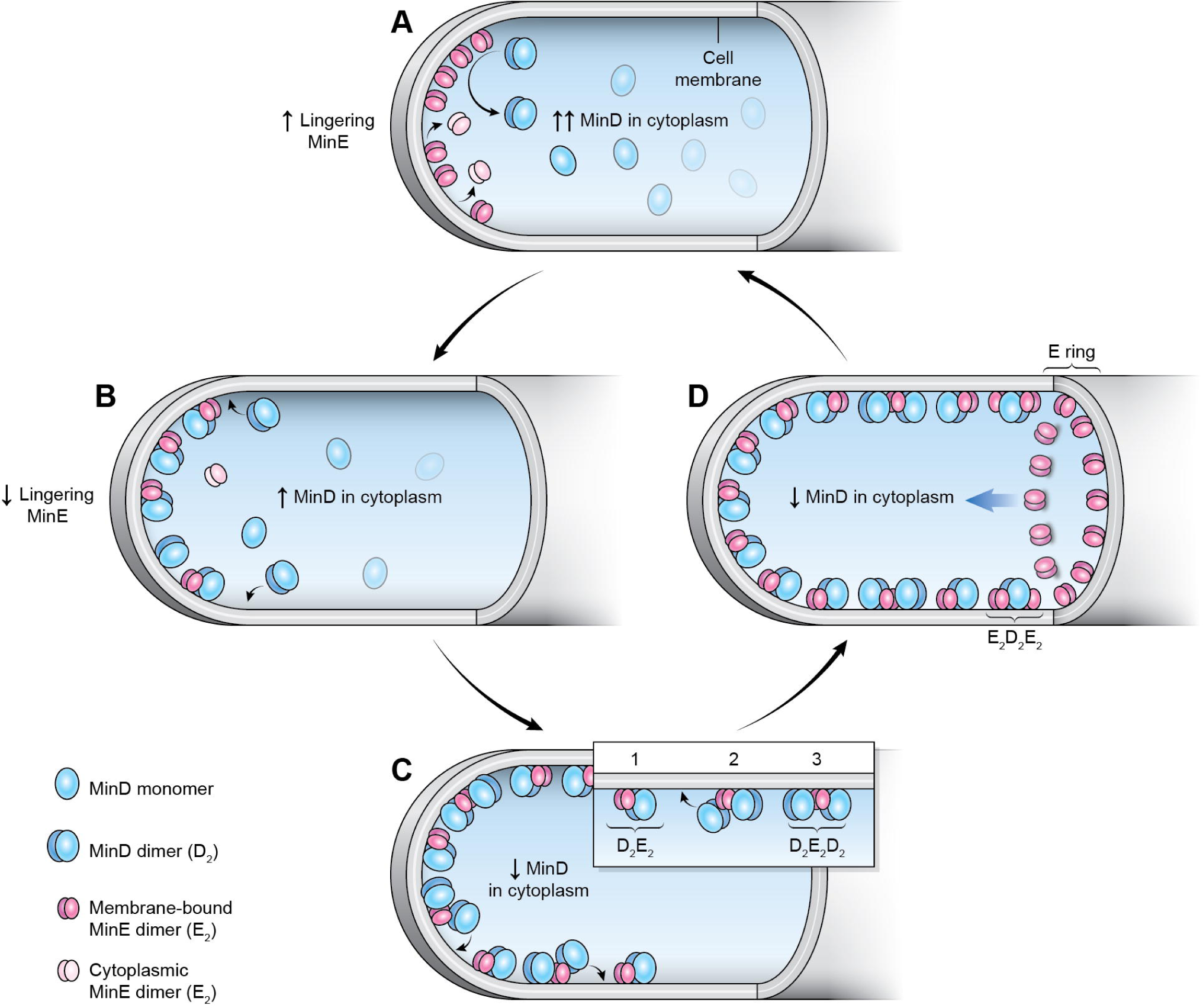
**Molecular model of Min oscillation *in vivo*.** (**A**) At high density, lingering MinE provides the refractory period for oscillation by inhibiting MinD from rebinding the cell-pole from which it just dissociated. (**B**) Once the lingering MinE density declines, MinD dimers can bind the membrane and join MinE dimers in a one-to-one complex. (**C**) This D2E2 complex then stimulates the recruitment of more MinD, to form D2E2D2. (**D**) As MinD depletes from the cytoplasm and MinE accumulates on the membrane, E2D2E2 can form, where a MinD dimer is now sandwiched by 2 MinE dimers. This complex stimulates MinD ATPase activity and release, allowing lingering MinE to concentrate into an E-ring. And the process repeats.

Here, we provide evidence that furthers our molecular mechanism for Min patterning; with an emphasis on how the multiple states of MinE drive oscillation by spatiotemporally regulating MinD interaction with the membrane.

## RESULTS AND DISCUSSION

### MinE dynamically associates with membrane independent of MinD

Structural studies of MinE alone, or in complex with MinD, have unveiled its conformational plasticity (Ghasriani et al., 2010; Kang et al., 2010; Park et al., 2011). MinE dimers are considered to be ‘closed’ or ‘inactive’ in solution and presumably in the cytosol *in vivo*. The MTSs of an ‘inactive’ dimer are packed against a six-stranded β-sheet at the dimer interface stabilizing the hydrophobic core (Figure 4A). The MinD-binding domains, which in their active state form α-helices adjacent to the MTSs, are occluded as the inner-most pair of β-strands at the dimer interface. Thus, the membrane and MinD interaction interfaces are for the most part unavailable or ‘closed’.

**Figure 4.**
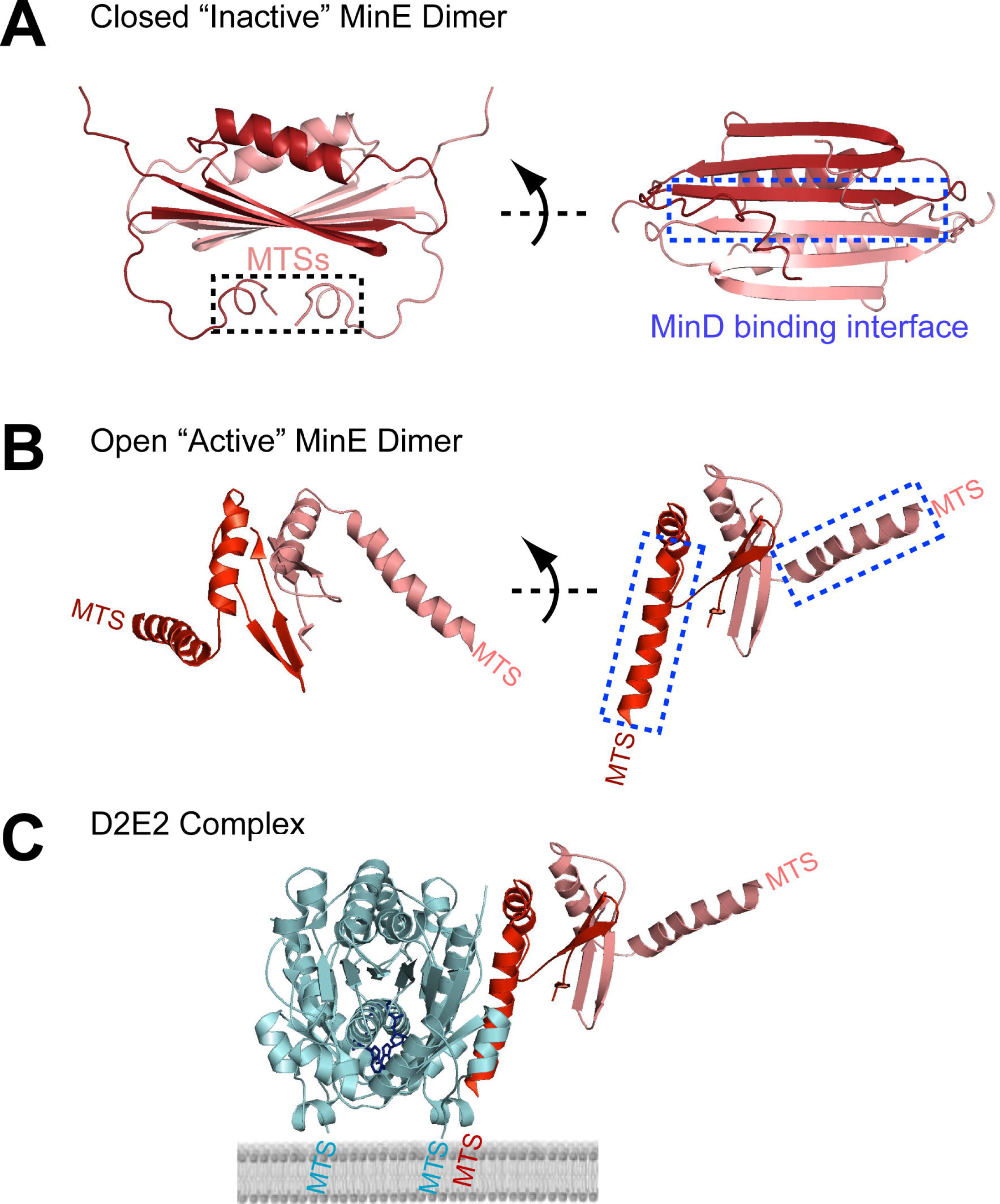
**The conformational gymnastics of MinE.** (**A**) The ‘inactive’ or closedstructure of the MinE dimer. MinD binding domains are in a non-binding conformation comprising the dimer interface as well as the loops that connect to the adjacent Membrane Targeting Sequences (MTSs). The MTSs are tacked onto this hydrophobic core. Structures adapted from PDB ID 2KXO (Ghasriani et al., 2010). (**B**) The ‘active’ or open structure of the MinE dimer in its MinD-interacting conformation. The once buried MinD binding interfaces and adjacent MTSs are now accessible for interaction. The MinD-interaction domains are likely in a random coil conformation when not bound to MinD. (**C**) The D2E2 complex. The open form of the MinE dimer (red) is stabilized upon interaction with the membrane-bound MinD dimer (cyan). The D2E2 complex is stably bound to the membrane via three MTSs, two from MinD and one from MinE. Structures for (B-C) adapted from PDB ID 3R9J (Park et al., 2011) for conceptual illustration purpose only. The MTS domains of MinD and MinE were not included in the PDB structures.

How then does MinE become ‘open’ and ‘active’? When flowing 1.5 μM MinE together with 1 μM MinD onto SLBs composed of *E. coli* lipid extract, or synthetic lipid mixtures with anionic lipid densities similar to that of *E.coli* membrane, the MinE density on the SLB prior to pattern formation was not zero (Ivanov and Mizuuchi, 2010; Vecchiarelli et al., 2014; Vecchiarelli et al., 2016). In buffers of lower ionic strength, or if the bilayer had a high content of anionic lipid, membrane binding by MinE was significant, even without MinD (Hsieh et al., 2010; Vecchiarelli et al., 2014). From this we proposed that inactive MinE in solution is in equilibrium with a small proportion of MinE dimers that are active for membrane binding.

It is attractive to speculate that the MTSs of a closed MinE dimer are transiently accessible for membrane interaction. If the association occurs next to a membrane-bound MinD dimer, the accessible portion of the MinD-binding interface on MinE could begin to refold into the MinD-interacting α -helix. The subsequent release of the innermost pair of β-strands at the dimer interface would complete the refolding of the entire MinD-interacting α-helix (Figure 4B). The resulting D2E2 complex would stabilize both MinD and MinE dimers on the membrane (Figure 4C). We propose that conformational fluctuations within the ‘inactive’ MinE dimer is coupled to transient interaction with membrane, which plays a pivotal role in both the long range inhibition, and short range nucleation, of membrane binding by MinD.

The idea of MinE membrane binding prior to pattern initiation was based on experiments in the presence of MinD. To directly test the membrane binding activity of MinE alone, we measured the SLB-bound equilibrium density of MinE dimers in the absence of MinD. With 1.5 μM MinE (monomer) in solution, 82 +/- 25 MinE dimers were bound per μm^2^ of the SLB, and FRAP was substantially faster than the one second time resolution of our microscope. The data show that MinE dimers can indeed bind membrane with fast exchange and independent of MinD.

### Patterns initiate by MinE locally stimulating MinD binding to the membrane

Previous models postulate that pole-to-pole oscillations result from MinD first binding the cell-pole *via* an enigmatic auto-catalytic process as first postulated by Meinhardt and de Boer, 2001. The models state that membrane-bound MinD recruits additional MinD from solution, inferring that MinD dimerization or some higher-order oligomerization largely takes place on the membrane. To directly test whether MinD alone can indeed bind the membrane in an auto-catalytic manner, we preincubated GFP-MinD (1 μM) with ATP to generate dimers competent for membrane binding. The sample was then flowed at 1 μl/min into the SLB-coated flowcell. MinD slowly and uniformly bound the SLB without any sign of auto-catalytic binding from a nucleation center, or rate acceleration after the initial binding event (Figure 5A, Movie 1). Rather, a homogeneous steady state MinD density was achieved after ~ 45 minutes of constant flow. We conclude that when MinD-ATP binds membrane on its own, there is no notable positive feedback operating to nucleate a MinD binding center.

**Figure 5.**
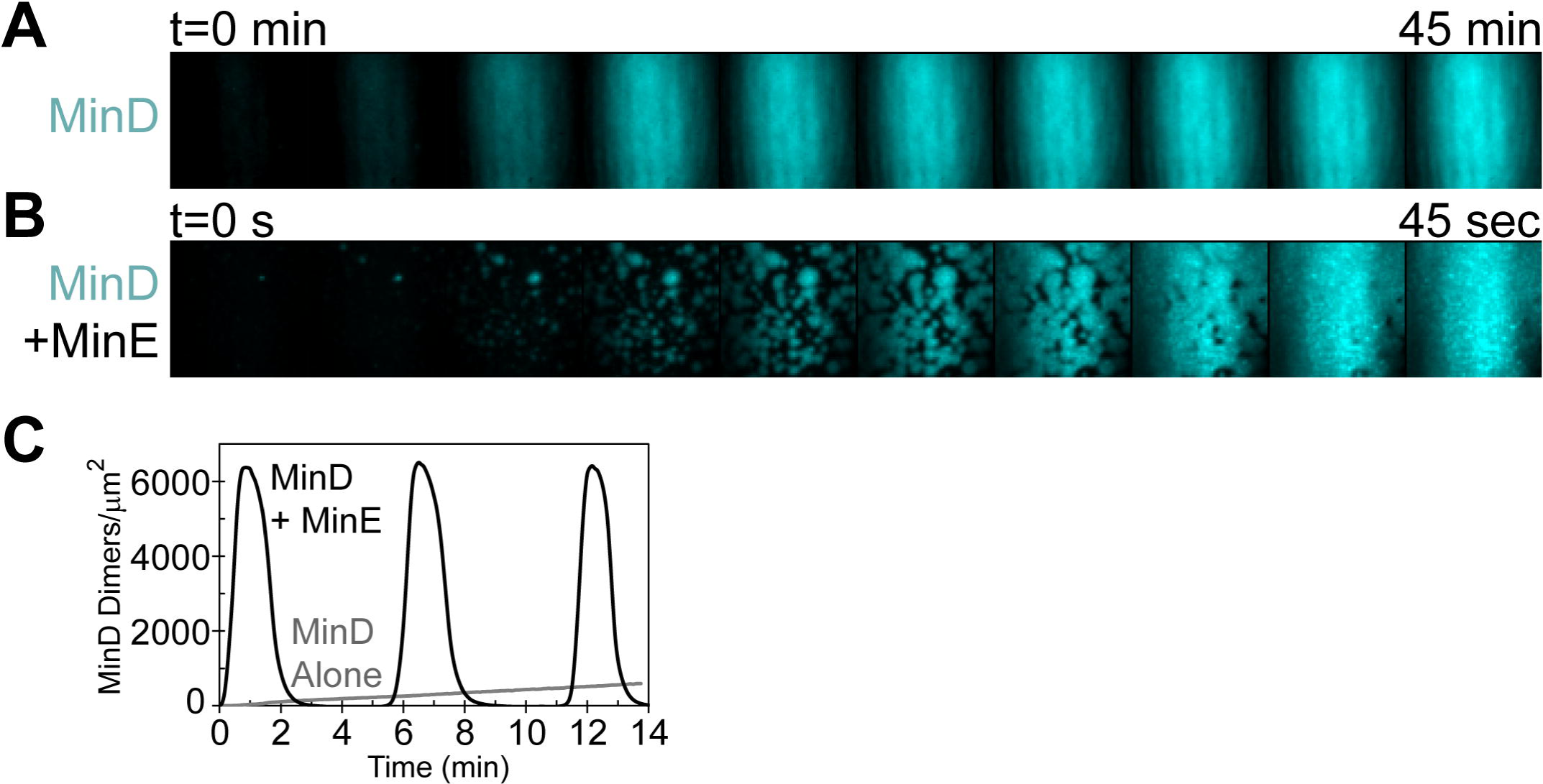
**MinE stimulates membrane binding by MinD** (**A**) MinD-ATP (cyan) homogeneously binds the SLB and slowly accumulates up to a maximum density. GFP-MinD (1 μM) in a buffer containing 2.5 mM ATP was flowed at 1 μl/min into an SLB-coated flowcell made of *E.coli* lipid. (**B**) In the presence of 5 μM MinE, MinD binding occurred rapidly from the radial expansion of nucleation points. The radially expanding zones merge as MinD accumulated up to the maximum density. (**C**) Quantification of MinD density on the SLB over time with or without MinE. The grey line shows the slow accumulation of MinD-ATP in the absence of MinE as shown in (A). At the same flow rate, MinE supports oscillation with MinD on the SLB (black line) as shown in (B). See Movie 1.

In stark contrast, when 5 μM MinE was also present in the sample, a short period of low binding by MinD and MinE was followed by the nucleation of rapid MinD binding from points on the SLB (Figure 5B, Movie 1). The radially expanding binding zones quickly merged and MinD reached maximum binding in under a minute. Under this constant sample flow, MinD and MinE created a spatially near-homogeneous oscillation on the SLB (Figure 5C, Movie 1) as previously described (Ivanov and Mizuuchi, 2010, Vecchiarelli et al., 2014). The data support the idea that MinE is required to nucleate and stimulate the rapid radial expansion of MinD polar zones *in vivo*.

We propose that the initial low-level binding of MinD and MinE on the bilayer and the local fluctuations of their relative ratio play key roles on when and where membrane binding by MinD is nucleated by MinE. The incredibly fast membrane binding equilibrium of MinE without MinD, as measured by FRAP, indicates that MinE binding to the membrane is faster than MinD at the start of our experiments. In this case, MinD-ATP dimers landing on the SLB would encounter membrane-bound MinE to form D2E2. At the initially low ratio of MinD to MinE on the bilayer, D2E2 would soon encounter another membrane-bound MinE dimer to become E2D2E2, which releases MinD from the membrane. In this pseudo-steady state, the surface density of MinE remains roughly constant while MinD dimers that attempt to bind the bilayer are quickly turned over. We propose this represents the lag phase that we observe prior to pattern formation in our cell-free setup (Ivanov and Mizuuchi, 2010; Vecchiarelli et al., 2014; Vecchiarelli et al., 2016).

The surface densities of MinD and MinE prior to pattern initiation are low, but highly responsive to local fluctuations in their ratio. According to our model, D2E2 can also recruit a MinD dimer to the membrane faster than MinD binding on its own. The probability of this happening would increase when D2E2 collisions with another MinE dimer on the membrane are delayed and when the concentration of active MinD dimers near the membrane increases. Thus, at a critical local state, the probability becomes significant enough for the balance to locally tip, which allows for MinE-stimulated nucleation of membrane binding by MinD. Once a MinD binding zone is locally nucleated, it radially expands, recruiting more MinE to form D2E2, which further stimulates the recruitment of more MinD to the membrane and so on. Therefore, the MinE-stimulated binding rate of MinD accelerates and converts all pre-existing lingering MinE dimers into the D2E2 complex, neutralizing their counteracting ability to dissociate MinD from the membrane. Thus, the switch from *preventing* to *stimulating* MinD binding is essentially determined by the balance of the membrane-bound MinE density and the solution concentration of active MinD-ATP dimers near the membrane.

### Lingering MinE provides the refractory period for oscillation by inhibiting MinD binding to membrane

Lingering MinE dimers are those remaining on the membrane after E2D2E2 triggers ATP hydrolysis, which causes MinD to release. Lingering MinE is still in its open and active form - competent for joining membrane-bound D2 to form D2E2, or with D2E2 to form E2D2E2. Our data has shown that the D2E2 complex binds membrane more stably than a MinD dimer on its own (Vecchiarelli et al., 2016). Lingering MinE would quickly join a MinD dimer on the membrane to form D2E2. Once the majority of membrane-bound MinD exists in this complex, any lingering MinE dimers remaining in the vicinity can transiently amplify themselves by disassembling D2E2 via E2D2E2 formation, resulting in a MinD dissociation feedback loop. Only when the D2E2 density on the membrane declines through this process, the level of lingering MinE can also diminish as it reverts to its inactive state and releases into solution on a time scale of several seconds. But as described earlier, instead of completely disappearing from the membrane surface, lingering MinE dimers diminish towards a low steady state level where the local MinD to MinE ratio on the membrane is highly responsive to the recovering active MinD concentration in solution that triggers the next nucleation event.

Lingering MinE is not restricted to 2D diffusion on the membrane. Rather, it can also diffuse briefly in solution. This active form of MinE in solution likely reverts back to the inactive state within a second or so, thus limiting its bulk diffusion distance. Solution diffusion of this MinE species explains our previous observation of circular MinD binding zones breaking symmetry to form waves that only travel upstream during sample flow (Vecchiarelli et al., 2014). Sample flow pushes the diffusible lingering MinE species downstream, thus only permitting upstream MinD binding and wave propagation.

Even in the absence of flow, it is likely that lingering MinE, transiently diffusing in solution, acts as a mesoscale spatial communicator for Min patterning. When a propagating wave approaches an amoeba for example, the wave seems to sense the E-ring of the amoeba at a distance, which then deforms the wave front (Figure 6A and Movie 3).

**Figure 6.**
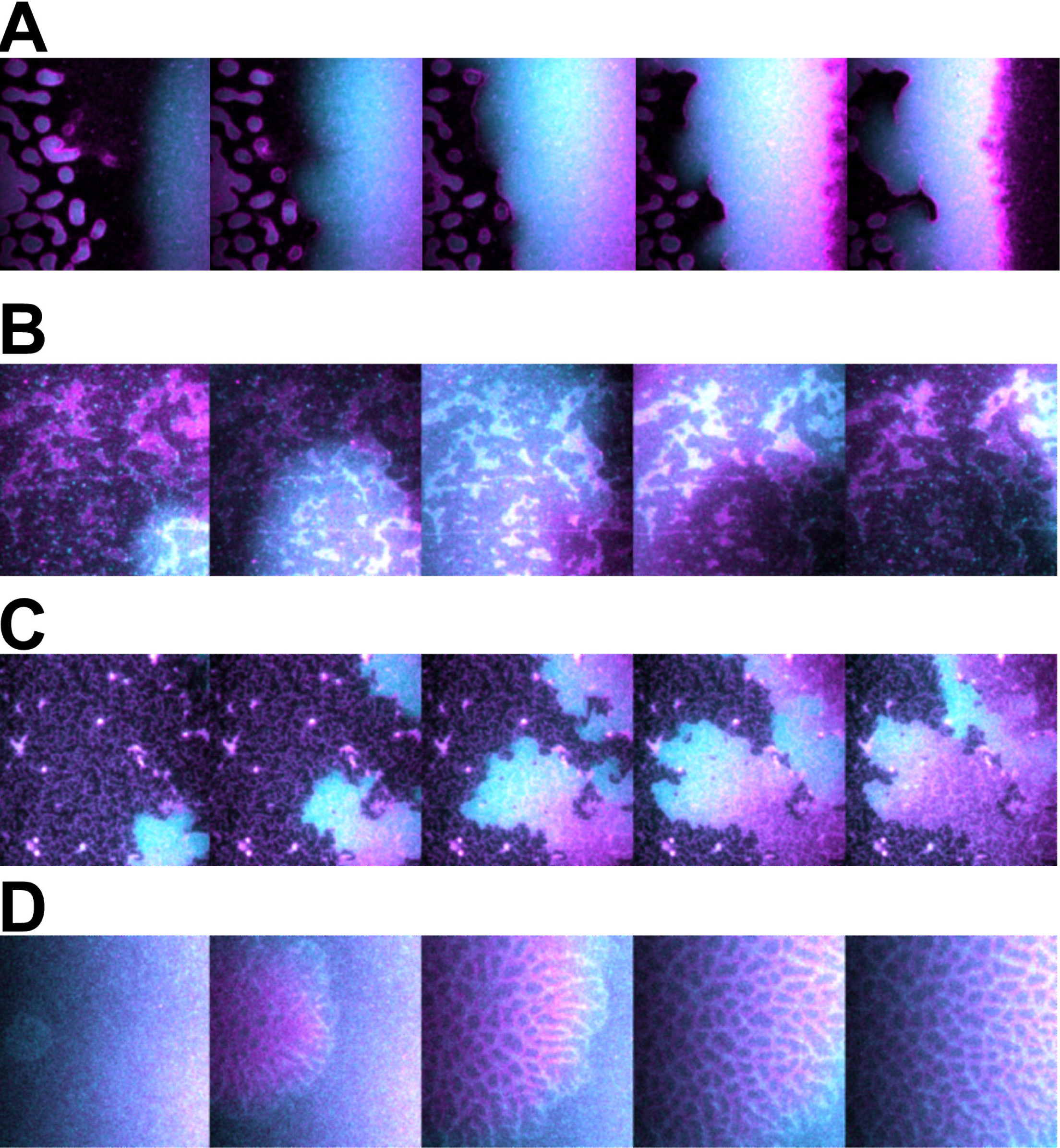
**How does MinE form an E-ring?** (**A**) Lingering MinE is a mesoscale spatial communicator for Min patterning. A freeze frame time course shows how a group of amoebas are approached by a travelling wave and disassembled at a distance. MinE within the E-ring of the disassembling amoebas seems to adsorb to the wave front several microns away on the SLB, resulting in a deformation of the wave. (**B**) Transitions in membrane structure alter Min patterning. A freeze frame time course of waves passing unhindered over patches of altered membrane state. (**C**) A freeze frame time course showing an example of where meshwork membrane transitions can become formidable barriers to wave propagation. (**D**) Freeze frame time course showing the nucleation and expansion of phase transitions in the SLB, as marked by MinE forming a stable meshwork around MinD-cores.

Spatial communication among patterns by this diffusible state of lingering MinE can also explain the previously reported wave phase synchronization across membrane gaps (Schweizer et al., 2012).

### MinE binding can cause membrane transitions that promote E-ring condensation

A big question remaining to be experimentally addressed is how does MinE coalesce into a high density and well-defined E-ring that resists diffusion? Our observations of Min patterning on less fluid SLBs provide insight. We have observed unhindered wave propagation with high density patches of MinD and MinE that form, fade, and reappear in the exact same position on an SLB (Figure 6B and Movie 4). But over time, some regions become formidable barriers to wave propagation (Figure 6C). We have also observed the nucleation of phase transitions in the SLB, as marked by MinE forming a stable meshwork around MinD-cores (Figure 6D). We previously found similar meshworks acting as barriers for wave progression (Ivanov and Mizuuchi, 2010).

What maintains this positional memory? We suggest these observations reflect a local transition in membrane structure and/or composition caused by MinE binding, which then acts as a positive feedback loop for further recruitment and stabilization of MinE on the membrane. These ‘frozen’ membrane patches were not as readily detectable when using synthetic lipid mixtures that maintain fluidity. The frozen mesh patterns we have observed could be exacerbated manifestations of MinE’s ability to condense into a thin tight E-ring without significant diffusional spreading by inducing local changes in membrane state (Vecchiarelli et al., 2016). MinE binding to membrane likely promotes dynamic local phase separations in membrane structure and/or composition. The nature of these membrane transitions remains to be further investigated and could shed light on how MinE dimers cooperatively form an E-ring without polymerizing into a filament.

### Travelling waves versus standing-wave bursts

Our recent findings have allowed us to reexamine the mechanism by which the Min system creates waves or bursts on an SLB *in vitro* (Vecchiarelli et al., 2016). Travelling waves were the first and most stable pattern formed on an SLB (Figure 7A) (Loose et al., 2008). Above waves, the amount of MinD and MinE molecules in solution relative to the membrane surface area are well in excess of that found *in vivo* (Shih et al., 2002) and the solution phase protein distribution is essentially homogeneous (Ivanov and Mizuuchi, 2010; Loose et al., 2008; Vecchiarelli et al., 2016; Vecchiarelli et al., 2014). When increasing the MinE concentration in solution, while keeping MinD constant, we found a corresponding increase in the rate of MinD binding at the wave front (Figure 7B). The data once again shows that MinE can stimulate MinD recruitment to the membrane.

**Figure 7.**
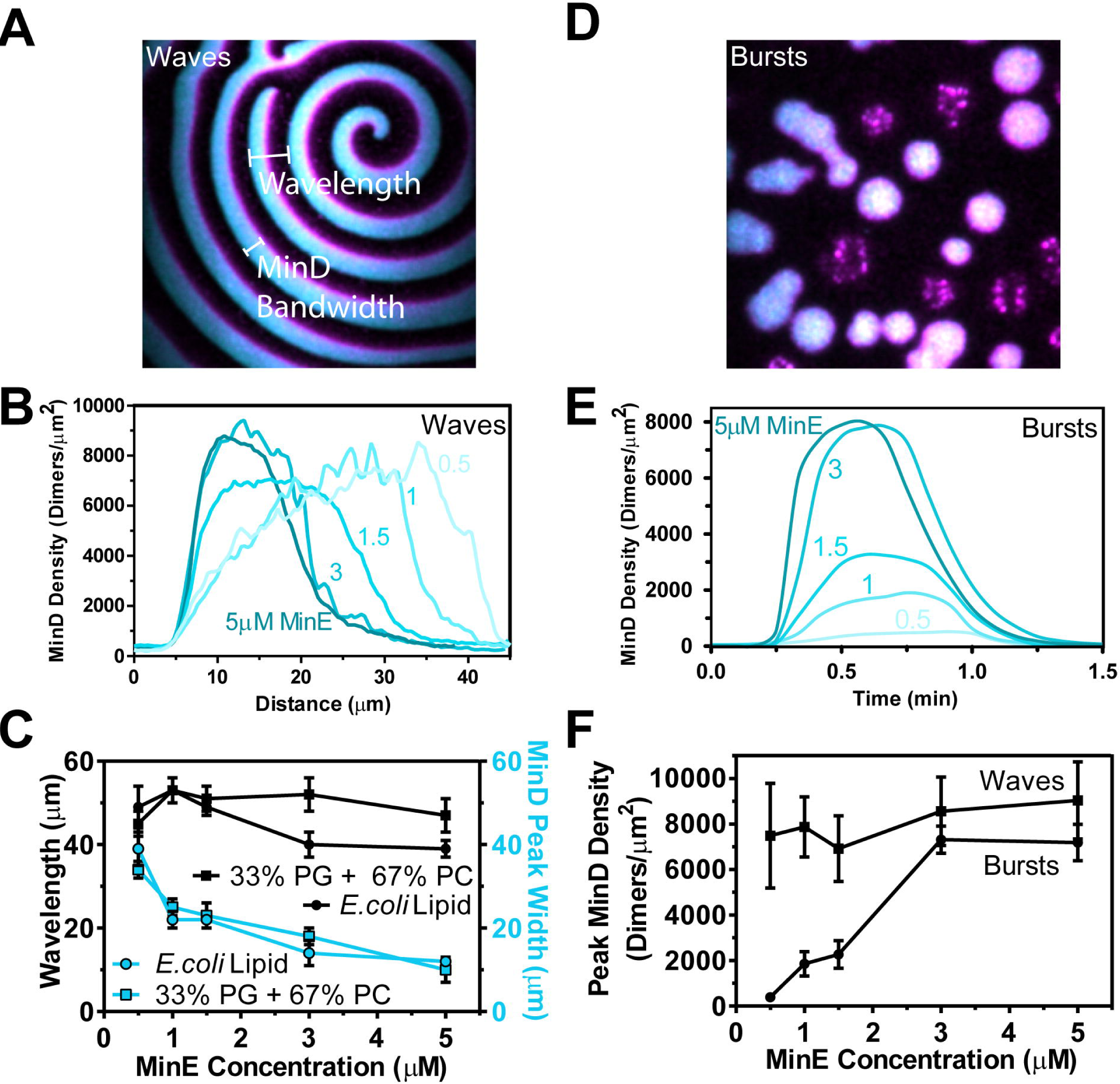
**Traveling waves versus standing-wave bursts.** (**A**) Freeze frame image of Min waves. The wavelength is the distance from one wave front to the next. MinD “peak-width” is the distance from the wave front to the MinE-rich rear where MinD has dissociated. (**B**) The MinD peak-width broadens as the solution concentration of MinE decreased. (**C**) Although MinD peak-width narrows with increasing MinE in solution, the wavelength remains constant. The MinD peak-width data were reproduced from Vecchiarelli et al., 2014 Figure 2E for comparison with the wavelength measurements. (**D**) Freeze frame image of Min bursts. (**E**) The rates of burst expansion and dissipation both increase with increasing MinE concentration. As a result, the temporal periodicity did not change. (**F**) Regardless of MinE concentration, waves have a saturating protein density on the SLB, which is responsible for the transition to protein release at the rear of a wave. Bursts on the other hand dissipate due to solution depletion of MinD.

Although higher MinE led to faster MinD binding at the wave front, the rate of MinD release at the rear did not dramatically change. Why does more MinE in solution not increase the release rate at the rear of waves as we have observed for bursts? The answer lies in the fact that, although more MinE in solution accumulates more MinD on membrane, the peak protein density of MinD achieved within a wave is essentially the same (Figure 7B), 8000 ± 2000 MinD dimers/μm^2^ (20-30% of surface confluence). Since the peak MinE density also reaches a similar level with lower MinE concentrations, MinE-stimulated disassembly of MinD at the rear of a wave occurs at a similar rate.

To summarize, more MinE in solution increases the rate at which MinD forms the wave front. But the peak MinD density remains the same. Although MinD density remains constant, MinE accumulates faster with higher MinE in solution, thus reaching the peak MinE density quicker and starting disassembly earlier. As a result, more MinE in solution narrows the MinD peak-width of a wave, but the wavelength itself does not change; 46 ± 6 μm on an *E. coli* SLB and 49 ± 4 μm on an SLB composed of 67% phosphotidylcholine (DOPC) and 33% phosphatidylglycerol (DOPG) (Figure 7C). The findings are consistent with the proposal of membrane-associated MinE being a catalyst for both membrane binding and release by MinD regardless of the pattern type.

Bursts form when the supply of active MinD is limiting (Figure 7D) (Vecchiarelli et al., 2016). Like waves, increasing the MinE concentration resulted in an increased rate of MinD binding during burst initiation and expansion, and the periodicity remained constant (Figure 7E). Unlike waves however, the peak MinD density within bursts increased with higher MinE in solution. Also, bursts were supported by protein densities significantly lower than those found in waves, due to depletion of the active MinD supply in solution (Figure 7F). As MinD binding slows, MinE binding catches up and disassembles the burst.

We conclude that bursts undergoing a standing-wave oscillation switch to disassembly due to the local depletion of active MinD in solution. For travelling waves, MinD depletion does not set the limit for MinD density on the SLB. Rather, we believe that protein-membrane interactions become strongly inhibited by surface exclusion. Consistently, the peak MinD protein density in waves does not exceed 20-30% of the surface density at confluence. At this density, the binding rate is expected to become very low due to surface area exclusion effects, deviating from predictions based on the Langmuir adsorption model (Schaaf and Talbot, 1989; Talbot et al., 1994). As the surface densities of D2 and D2E2 become higher than ~ 10% confluence, the rate of MinD binding will slow faster than MinE because MinE has a smaller footprint on the SLB. Once D2E2 becomes the more prevalent complex on the SLB, E2D2E2 would eventually form and start dissociating MinD from the SLB, making even more room for MinE binding. This scenario conveniently explains the plateau and decline of MinD that is accompanied by a transient acceleration of MinE binding just before the MinD to MinE ratio switches (see Ivanov and Mizuuchi, 2010). To put it simply, MinD stops binding within a wave because there is no more room on the SLB, whereas in bursts (or in a MinD polar zone *in vivo*), MinD stops binding because the solution (or cytoplasmic) supply has been depleted.

## CONCLUSIONS

A large body of cell-free observations has allowed us to propose a comprehensive molecular mechanism that explains the wide variety of patterns achievable by MinD and MinE self-organization on a membrane (Vecchiarelli et al., 2016). Although only two proteins are required for patterning, this ‘simple’ system actually involves a large number of key molecular species. Contrary to previous models showing MinD binding to the membrane in an auto-cooperative process and MinE coming along to kick it off, we find that MinE orchestrates the entire oscillatory process through regulation of MinD membrane binding. We conclude that MinE successively recruits, stabilizes, releases and inhibits MinD interactions with membrane to drive oscillation.

Based on the limited experimental data currently available, only a subset of reaction parameters have been estimated directly for mathematical modeling and simulation. A systematic biochemical study and quantitative analysis of each reaction step is essential to confirm several aspects of our proposed mechanism as well as to impose constraints on the rate parameters involved. These experimental approaches, combined with quantitative simulations, will further refine and improve our understanding of this fascinating and beautiful system.

## EXPERIMENTAL PROCEDURES

**Proteins.** Protein expression, purification, and fluorescent labeling were performed as previously described (Vecchiarelli et al., 2014).

**Flowcell Assembly.** Flowcell assembly (Vecchiarelli et al., 2015) and bilayer coating with E. coli polar lipid extract or monounsaturated (18:1) synthetic lipids (Vecchiarelli et al., 2014) has been previously described. The synthetic lipid mixture was composed of 67% 1,2-dioleoyl-sn-glycero-3-phosphocholine (catalog no. 850375), and 33% 1,2-dioleoyl-sn-glycero-3-[phosphor-rac-(1-glycerol)] (catalog no. 840475). All lipids were purchased from Avanti in chloroform at 25 mg/mL.

**Sample Handling and Preparation**. Experiments were performed in Min buffer: 25 mM Tris-HCl, pH 7.4, 150 mM KCl, 5 mM MgCl_2_, 2 mM DTT, and 0.5 μg/mL ascorbate.

Five millimolar phosphoenolpyruvate (Sigma) and 10 μg/mL pyruvate kinase (Sigma) provided ATP regeneration.

To form the Min pattern spectrum, His_6_-eGFP-MinD was mixed with MinE-His_6_ (mixed 1:19 with MinE-Alexa 647) at the concentrations specified and preincubated in Min buffer for 15 min at 23 °C before addition of 2.5 mM ATP in a final reaction volume of 500 μL. The sample was passed through a 0.2-μm Amicon filter and loaded into a 1-mL syringe. TFZL tubing (1/16 × 0.02 inch; UpChurch) connected the syringe to the flowcell inlet Nanoport (UpChurch). Samples were infused into the 3-μL flowcell (25 μm × 4 mm × 30 mm) with a neMESYS pump (Cetoni) at 1 μL/min (cross-sectional average velocity of 0.17 mm/s) for 10 min. Flow was stopped before movie acquisition, except for the MinD membrane binding assay where flow maintained during movie acquisition.

**MinD-independent MinE binding to an SLB.** To convert fluorescence intensity to an estimate of MinE dimers on the SLB as well as to measure the background fluorescence contributed by the solution concentration of MinE in the evanescent illumination volume, 2 or 4 μM MinE-Alexa488 (~ 60% labeled) was mixed in reaction buffer without ATP and flowed at 5 μl/min onto an SLB composed of DOPC alone, which MinE does not bind. Fluorescence intensity was measured before flowing in MinE to establish background. Then the MinE sample was flowed into the flowcell, and then washed away with buffer to ensure the intensity fell to background levels. The fluorescence signal of MinE in the solution volume within the evanescence illumination depth was obtained by subtracting the background signal with buffer only (mostly camera dark noise). From the wavelength, the refractive indices of fused silica and the reaction buffer, and the illumination angle, we calculated the evanescence penetration depth to be 131 nm. From the known sample concentrations of MinE, the detected fluorescent signal was then converted to the number of MinE dimers in the evanescent volume. The same experiments were then carried out on our standard SLB composed of 67% phosphotidylcholine (DOPC) and 33% phosphatidylglycerol (DOPG) that MinE binds. From the fluorescence signal, the background contribution was subtracted, which included MinE fluorescence in solution. The background-subtracted fluorescence signal attributable to SLB-bound MinE dimers was then converted to a molecular density per unit surface area, using the conversion factor calculated above.

**Total Internal Reflection Fluorescence Microscopy**. Total Internal Reflection Fluorescence (TIRF) illumination and microscopy as well as camera settings were as previously described (Vecchiarelli et al., 2014). Prism-type TIRF microscopy was used with an Eclipse TE2000E microscope (Nikon) with a PlanApo 10× (N.A. = 0.45, air) or 40× (N.A. = 1.0, oil-immersed) objective lens. The TIRF illumination had a Gaussian shape in the field of view; therefore, intensity data for Min protein density estimations were measured at or near the middle of the illumination profile.

Movies were acquired using Metamorph 7 (Molecular Devices) and transferred to ImageJ (National Institutes of Health) for analysis and conversion to AVI file format. Brightness and contrast were set for each image or movie acquisition individually to best represent the features discussed. However, paneled acquisitions in the same movie share the same settings. All data were acquired at 5 s/frame when using full-length MinE. When using MinE_11–88_, the frame rate was 1 s/frame. Accelerations are indicated in the movie legends.

**TIRF Microscopy Protein Density Estimation** during pattern formation. The average fluorescence intensity of single GFP-MinD or MinE-Alexa647 molecules was measured as previously described (Vecchiarelli et al., 2014) to calculate the Min protein density on the SLB expressed as dimers/μm^2^.

## ACKNOWLEDGMENTS

This work was supported by the intramural research fund for NIDDK (to K.M.), NIH US Dept of HHS, and the Nancy Nossal Fellowship (to A.G.V.).

### AUTHOR CONTRIBUTIONS

A.G.V. and K.M. designed the study. A.G.V. performed the majority of experiments described. M.L. and M.M. did the MinE-membrane interaction experiments. V.I. did the membrane transition experiments. A.G.V. and K.M. wrote the paper.

## MOVIE LEGENDS

**Movie 1. MinE stimulates membrane binding by MinD**. The left panel shows MinD-ATP (cyan) slowly and homogeneously binding the SLB. GFP-MinD (1 μM) in a buffer containing 2.5 mM ATP was flowed at 1 μl/min into an SLB-coated flowcell made of *E.coli* lipid. The movie on the right shows that in the presence of MinE (magenta), MinD (cyan) binding occurs more rapidly from the radial expansion of nucleation points that merge with constant flow. This is followed by the rapid disassembly of both proteins. Constant flow supports a near spatially homogeneous oscillation. Movies are 100 times faster than real time. Panel areas are 100 × 100 μm. Related to Figure 5.

**Movie 2. Lingering MinE is a mesoscale spatial communicator for patterning**.

Lingering MinE (magenta) from the E-rings around the MinD (cyan) cores of amoebas are transferred at a distance to an approaching wave front. Movies are 100 times faster than real time. Panel areas are 100 × 100 μm. Related to Figure 6A.

**Movie 3. Examples of altered Min patterning on bilayers of compromised integrity.**

The left panel shows waves passing unhindered over patchy regions of the SLB. The middle panel shows how membrane transitions can become formidable barriers to wave propagation. The right panel shows the nucleation of a phase transition in the SLB, as marked by MinD and MinE forming a stable meshwork. Movies are 100 times faster than real time. Panel areas are 100 × 100 μm. Related to Figure 6B-D.

